# Rapid Recalibration of Peri-Personal Space; Psychophysical, Electrophysiological, and Neural Network Modeling Evidence

**DOI:** 10.1101/842690

**Authors:** Jean-Paul Noel, Tommaso Bertoni, Emily Terrebonne, Elisa Pellencin, Bruno Herbelin, Carissa Cascio, Olaf Blanke, Elisa Magosso, Mark T. Wallace, Andrea Serino

## Abstract

Interactions between individuals and the environment are mediated by the body and occur within the peri-personal space (PPS) – the space surrounding the body. The multisensory encoding of this space plastically adapts to different bodily constraints and stimuli features. However, these remapping effects have only been demonstrated on the time-scale of days, hours, or minutes. Yet, if PPS mediates human-environment interactions in an adaptive manner, its representation should be altered by sensory history on trial-to-trial timescale. Here we test this idea first via a visuo-tactile reaction time paradigm in augmented reality where participants are asked to respond as fast as possible to touch, as visual object approach them. Results demonstrate that reaction times to touch are facilitated as a function of visual proximity, and the sigmoidal function describing this facilitation shifts closer to the body if the immediately precedent trial had indexed a smaller visuo-tactile disparity (i.e., positive serial dependency). Next, we derive the electroencephalographic correlates of PPS and demonstrate that this measure is equally shaped by recent sensory history. Finally, we demonstrate that a validated neural network model of PPS is able to account for the present results via a simple Hebbian plasticity rule. The present findings suggest that PPS encoding remaps on a very rapid time-scale and is sensitive to recent sensory history.

## Introduction

Physical interactions between an agent and the environment happen by mediation of the body and occur within the peri-personal space (PPS; Rizzolatti et al., 1981) – the space immediately adjacent to and surrounding one’s body (Rizzolatti et al., 1997; di Pellegrino et al., 1997). This space is encoded by a dedicated fronto-parietal neural network, in which neurons possess visuo-tactile receptive fields anchored on particular body parts – most notably the face and hand, but also the trunk (Graziano et al., 1994, 1997, 1999; Duhamel et al., 1998; Schlack et al., 2005; see Serino et al., 2015b, for an analogous demonstration in humans). These multisensory neurons are most responsive to dynamic looming stimuli (Fogassi et al., 1996; Noel et al., 2018b), and are part of a network which can evoke complex and stereotyped defensive movements upon microstimulation (Cooke et al., 2003; Cooke & Graziano, 2004). In turn, PPS is conceived as an adaptive multisensory space mediating both bodily protection and goal-directed action (Cooke & Graziano, 2004; Brozzoli et al., 2014; Serino, 2019).

Studies show that the shape and size of PPS is not fixed, but instead adapts as a function of interaction with the environment both dynamically and plastically. For instance, visual-tactile coding of the peri-hand space modifies online – in a dynamic fashion – during planning (Patanè et al 2018) and execution (Brozzoli et al., 2009, 2010) of grasping actions. Further, PPS extends in space after few minutes of using a tool to reach far locations (Iriki et al., 1996; Maravita & Iriki, 2004; Farne & Ladavas, 2000; Canzoneri et al., 2013), or conversely it contracts after prolonged immobilization (Bassolino et al., 2015), or shifts to reflect altered self-location (Noel et al., 2015a, 2019; Salomon et al., 2017). Arguably, the largest portion of the effort in studying PPS today centers on delineating its remapping as a function of a number of external manipulations on the order of days, hours, or minutes (see references above).

However, if the encoding of the space immediately surrounding one’s body is truly fundamental in mediating sensorimotor affordances and interaction with potential threats, this encoding must update as the environment and sensory history changes. That is, in addition to the already demonstrated dynamic updating within trials (e.g., Fogassi et al., 1996; Noel et al., 2018) and slow plastic updating between trials (e.g., Canzoneri et al., 2013) there must be a third intermediate mode: *rapid* and plastic updating incorporating new sensory evidence. Classic adaptation-recalibration studies in multisensory temporal acuity demonstrated early on that upon extensive sensory exposure, perceptual judgments shifted as to reflect the statistics of the presented stimuli (Fujisaki et al., 2004; Vroomen et al., 2004; Noel et al., 2015c). More recently, these effects have been shown to occur on a trial-by-trial basis (positive serial dependency; Van der Burg et al., 2013, 2015; Noel et al., 2016a; 2016b). In analogy to these recent findings within the multisensory temporal domain, we asked whether rapid recalibration applies to the multisensory mechanisms underlying PPS. This putative occurrence would suggest that PPS can be regulated online to adapt to changes in the environment.

We present psychophysical (Experiment 1) and electrophysiological (Experiment 2) evidence for the rapid recalibration of PPS, as well as a neural network model implementing a neurophysiologically plausible mechanism for this recalibration (Experiment 3). In Experiment 1, we index rapid recalibration of PPS in a naturalistic context via a behavioral paradigm implemented in augmented-reality (Serino et al., 2017). Participants were presented with looming visual stimuli and at a given distance between the body and the visual stimuli, target tactile stimulation was delivered. Indexing PPS, reaction times to touch are expected to decrease as a function of decreasing visuo-tactile distances (Serino, 2019). Critically, if rapid recalibration occurs, here we expect the facilitation to touch as a function of visuo-tactile distance to be more pronounced (occur at larger spatial disparities) when the immediately precedent trial probed a larger, as opposed to smaller, visuo-tactile distance. In Experiment 2, we extend an intracortical local field measure of PPS (Bernasconi et al., 2018, see also Noel et al., 2018c, 2019a) to scalp electrophysiology (EEG), and demonstrate a PPS rapid recalibration correlate akin to that observed in multisensory temporal judgments (Simon et al., 2017). Lastly, to suggest a mechanistic account, we demonstrate that a validated neural network model of PPS (Magosso et al., 2010a,b) can in principle account for the rapid recalibration of PPS given Hebbian learning (see Serino et al., 2015a).

## Methods

### Experiment 1 - Psychophysics

#### Participants

Thirty-eight (mean age = 22.9 ± 0.86, range 19-44) right-handed students from the Ecole Polytechnique Federale de Lausanne took part in this experiment. This sample size is approximately twice as large as most behavioral studies of PPS (e.g., 20 participants in Noel et al., 2018a, b, e), given that we anticipated needing to discard a large portion of subjects due to the multiple data fitting procedures (3 separate sigmoidal fits as opposed to a singular one, see below). All participants reported normal touch and had normal or corrected-to-normal eyesight. All participants gave their written informed consent to take part in this study, which was approved by the local ethics committee, the Brain Mind Institute Ethics Committee for Human Behavioral Research at EPFL, and were reimbursed for their participation.

#### Materials & Apparatus

A Mixed/Augmented-reality technology (Figure 1A. Reality Substitution Machine – RealiSM; http://lnco.epfl.ch/realism) was utilized in order to render a pre-recorded panoramic physical scene (a looming ball originating at approximately 2 meters and traveling at a velocity of approximately 75 cm/s; recorded with 14 GoPro Hero4 cameras placed in a spherical rig - 3D 360hero 3DH3PRO14H) alongside the veritable surrounding environment (the experimental room) in which the participants were placed. Real-time stereoscopic images of the participant’s body were captured via a DuoCam (Duo3D MLX, 752×480 at 56Hz) and participant’s limbs were rendered online (delay < 5 ms). The visual stimulation was displayed on an Oculus Rift DK2 head-mounted display (HMD; 900×1080 per eye, 105° FOV). Tactile stimulation (duration; 6 ms up/down state) was administered via a solenoid (MSTC-3 tappers, M & E Solve).

**Figure 1.**
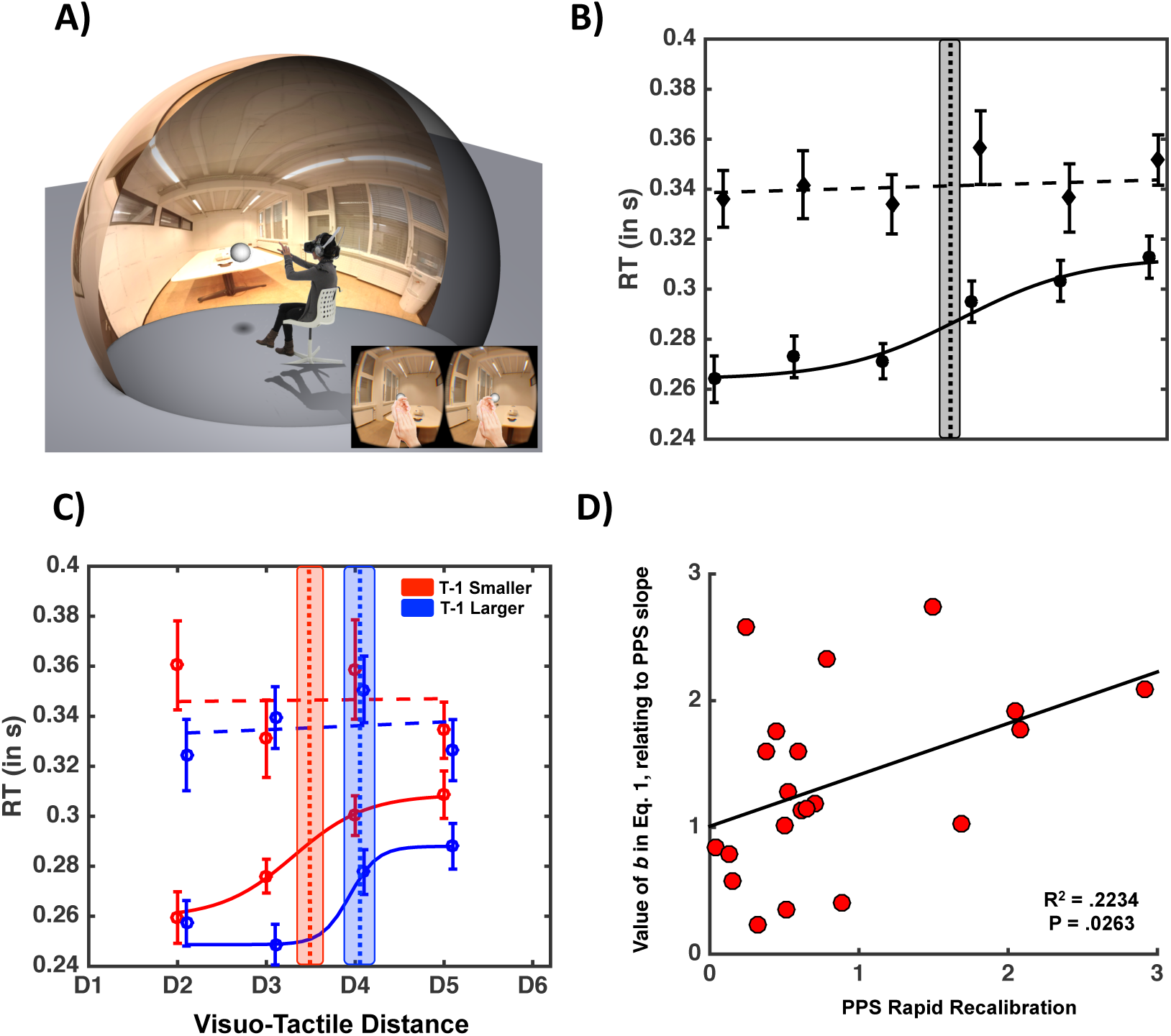
**A)** Experiment Setup. Participants are placed within an augmented-reality scenario in which they perceive a virtual looming ball approach them in their actual physical environment. Participants are equally equipped with a tactile tapper that provides somatosensory stimulation at a varying visuo-tactile distance. See text for further detail. **B)** PPS representation; the closer (D1 = closest, D6 = farthest) a task-irrelevant visual stimulus was from the participant the faster are RTs to touch (solid line: about a 40 ms facilitation). This effect cannot be explained as an expectancy effect, as baseline trials (dashed line) in which no visual stimulus was administered did not portray the same facilitation. Error bars represent +/- 1 S.E.M, vertical dashed line indicates average PPS boundary, and shaded area around it indicates +/- 1 S.E.M in central point estimate. **C)** PPS representation as a function of the immediately precedent trial; the sigmoidal curve indexing PPS is shifted toward the right (left), respectively, blue and red solid line, when trial t-1 probed a larger (smaller) PPS representation. Error bars, vertical dashed line, and shaded area around vertical line are as **B). D)** Relation between the parameter *b*, that contributes to defining the gradient of PPS, and rapid recalibration. Participant with a softer transition between the near and far space (as indexed by the value of parameter b) demonstrate a greater degree of rapid recalibration of their central point on a trial-per-trial basis (as indexed by the difference between their central point when T-1 was larger vs. smaller). The relation here is positive (as opposed to negative), as by Eq. 1., the larger the b value is, the flatter the slope of the sigmoidal. See Supplementary Material, Figure S1, for a similar analysis including all participants and without the fitting procedure.

#### Procedure

Participants were seated in a dimly lit room (see Figure 1A) in which they performed a tactile reaction time (RT) task to a stimulation administered on their right cheek. Each trial began with a white fixation cross, presented in the center of the virtual environment for 1.2s. Critically, in experimental trials (72% of all trials) they concurrently viewed a three-dimensional ball loom toward them in augmented reality (see Serino et al., 2015b, 2017 for a similar approach). This visual stimulus started approaching participants 300ms following fixation cross disappearance, and the onset of tactile stimulation was staggered with regard trial start by either T1 = 1.83s, T2 = 2.15s, T3 = 2.47s, T4 = 2.79s, T5 = 3.11s, or T6 = 3.43s. These temporal offsets map onto the spatial dimension linearly and negatively when stimuli approached the body at a constant velocity, and thus we denominate in the spatial dimension D1 = T6, D2 = T5, and so on. Therefore, the visuo-tactile disparities tested were D1 = 55.25 cm, D2 = 79.25 cm, D3 = 103.25cm, D4 = 127.25cm, D5 = 151.25cm, and D6 = 175.25cm.

In addition to the experimental trials we also included unisensory tactile trials (16% of all trials). These trials acted as a baseline condition in which tactile stimulation was administered at the equivalent temporal offsets as in experimental trials (T1 through T6), but no visual stimulus was presented. These trials are fundamental in ascertaining that putative effects in the experimental trials are veritably due to multisensory interactions, and not simply due to an expectancy effect (which would be time-dependent and thus revealed in the baseline trials). Lastly, catch trials in which visual stimuli were presented but no tactile stimulation was administered (12% of total trials) were undertaken in order to monitor task-compliance. All trial types were randomized within- and between-subjects and the inter-trial interval was set to 500ms. Every subject performed a total of 300 trials (36 repetitions X (6 experimental conditions + catch) + 8 repetitions X 6 baseline conditions).

#### Analysis

In a first step, participant’s RTs were collected and averaged as a function of condition (and regardless of the nature of the precedent trial). As a preliminary analysis, we performed a 2 (Condition: Experimental vs. Baseline) X 6 (Distance: D1 through D6) within-subject ANOVA in order to confirm that 1) multisensory visuo-tactile trials are faster than unisensory tactile trials, 2) visuo-tactile trials exhibit a space-dependence, but 3) tactile trials do not. Next, returning to the raw data we split experimental and baseline trials (t) on whether they had been preceded (t-1) by either a smaller or a larger visuo-tactile/tactile distance. That is, say trial t administered tactile stimulation when the visual stimulus was placed at D3. This trial was sorted into ‘D3, T-1 smaller’ if t-1 was a D1 or D2 trial, or ‘D3, T-1 larger’ if t-1 was a D4, D5, or D6 trial. Trials that were preceded by either the same distance (D3 preceded by D3) or a catch trial were omitted, as were D1 and D6 trials as, respectively, there was no smaller / bigger condition for these. Subsequently to this conditional sorting of trials by the nature of the immediately prior trial, on a subject-per-subject basis we fit the average RTs to a sigmoidal function (Eq. 1) from which we extract the central point of the sigmoidal (*x*_*c*_, in Eq. 1, representing the boundary of PPS) and a parameter proportional to its slope at the central point (*b* in Eq. 1, characterizing the gradient of PPS representation; See Canzoneri et al., 2012 for a similar approach). Participants were discarded from further analysis if one of their fits (i.e., regardless of sensory history, T-1 Smaller, or T-1 Larger than current) were dissatisfactory (a priori set to R^2^ < .50, 17 subjects removed. 21 subjects had R^2^ > 0.75 for all three experimental conditions). Central point and slope, were compared as a function of T-1 Smaller or Larger by means of a paired-samples t-test.

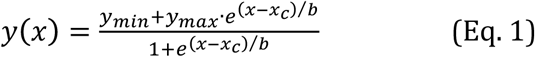

### Experiment 2 – Electroencephalography

#### Participants

Twenty-seven (15 females, mean age = 21.3 ± 0.79, range 18-31) right-handed students from Vanderbilt University took part in this experiment. All participants self-reported normal touch and had normal or corrected-to-normal eyesight. Six participants were excluded from analysis due to excessive motion and/or blink artifacts in the EEG recordings resulting in >50% of trials being rejected (3 subjects), large degrees of electrical noise (1 subject), technical problems (1 subject), and/or poor psychometric fits that precluded behavioral analysis (1 subject). Therefore, in total data from 21 participants (12 females) was analyzed. All participants gave their written informed consent to take part in this study, which was approved by the Behavioral Sciences Committee at Vanderbilt University.

#### Materials & Apparatus

The augmented reality setup from Experiment 1 was not used, due to the difficulty in recording high-density EEG while concurrently wearing a head-mounted display. Further, we decided to utilize static stimuli, as opposed to dynamically looming stimuli in Experiment 1 to render both visual and tactile stimuli ‘evoked’ (i.e., with sharp on-off transients), as opposed to having the tactile component be evoked, and the visual induced (i.e., prolonged presence and no transient visual on-off during tactile stimulation). Hence, visual and tactile stimuli were driven via a micro-controller (SparkFun Electronics, Redboard, Boulder CO) and direct serial port communication under the control of purpose written MATLAB (MathWorks, Natick MA) and Arduino (Arduino™) scripts. Visual stimuli were a flash of light presented by means for a red LED (3 mm diameter, 640 nm wavelength), while tactile stimuli consisted of vibrotactile stimulation administered via a mini motor disc (10mm diameter, 2.7mm thick, 0.9 gram, 5V, 11000 RPM). These stimuli were 50 ms in duration (square-wave, onset and offset <1 ms, as measured via oscilloscope). The LEDs and vibrotactile motor were mounted in an opaque enclosure where 30 LEDs sequentially protruded above the enclosure every 3.3 cm (in depth) and counted with a hand rest immediately adjacent to the first LED (Figure 2A, see Noel et al., 2018a for a similar apparatus). In the current study LEDs number 2, 5, 8, 11, 14, 17, and 20 were utilized, corresponding to visuotactile depth distances of 3.3cm, 13.2cm, 23.1cm, 33.0cm, 42.9cm, 52.8cm, and 62.9cm. Visuotactile stimuli consisted of the synchronous presentation of the visual and tactile stimuli described above.

**Figure 2.**
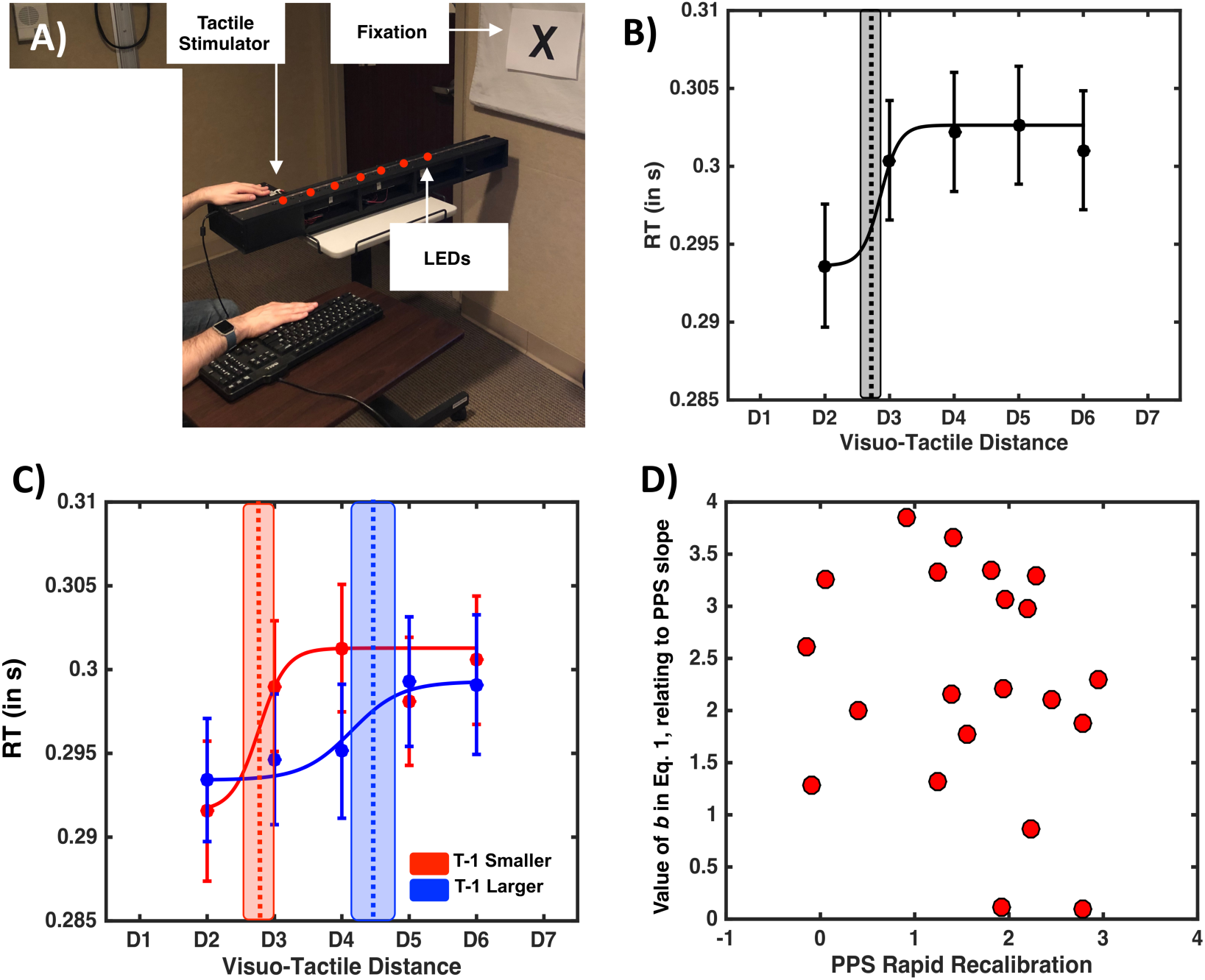
Behavioral Results. **A)** EEG setup. Participant’s viewed visual stimuli flash at different distances, while on given trials they simultaneously were given tactile stimulation on the hand. **B)** Tactile RT (y-axis) as a function of visuo-tactile distance (x-axis). Dots illustrate the group average RT, while error bars illustrate +/- 1 S.E.M. Dashed vertical line is the mean central point of the sigmoidal function describing tactile RTs as a function of visuo-tactile distance, and gray shaded area around the vertical line is the uncertainty (S.E.M) associated with the central point estimate. **C)** Tactile RT as a function of visuo-tactile distance (x-axis) and whether the precedent trial (i.e., t-1) was one presenting a larger (blue) or smaller (red) visuo-tactile spatial disparity. The rest of conventions as in left panel. **D)** Correlation between the degree to which a particular participant’s PPS recalibrated as a function of the previous trial (x-axis) and parameter *b* in Eq. 1 (y-axis) regardless of the nature of the previous trial.

#### Procedure

Participants were seated in a room with lights off (ambient light allowed for visibility), and performed a tactile reaction time (RT) task to stimulation administered on their left index finger. Participants were required to gaze toward a fixation point present at eye-level and placed at the end of the apparatus enclosing the LEDs (Figure 2A). Here, tactile stimuli was given on the index finger as opposed to the cheek – as in Experiment 1 – in order to minimize the potential for eye-blink artifact. Further, this change from Experiment 1 afforded us the possibility of exploring whether rapid recalibration of PPS applies generally to both peri-hand and peri-face space (and is present in both a veridical and augmented reality setting). Trials could be visuotactile (i.e., experimental trials; VT), tactile (i.e., baseline; T), or visual (i.e., catch; V) trials. Similar to Experiment 1, visual trials were ‘catch trials’ in that they did not require a motor response, while tactile trials were ‘baseline trials’ as these permitted us to gauge tactile RTs in the absence of visual inputs, and hence determine whether a multisensory effect was present as a function of visuo-tactile distance. Visuotactile trials were presented at 7 different distances (D1 through D7 = 3.3cm, 13.2cm, 23.1cm, 33.0cm, 42.9cm, 52.8cm, and 62.9cm; Figure 2A), while visual only trials were presented solely at D2, D3, D4, D5, and D6 due to time constraints and the fact that analyses heavily relied on distances between D2 and D6, as there were no smaller distances than D1 or larger distances than D7. Within each block 360 trials were presented; 40 VT trials at each of the 7 distances, 10 V trials at each of the 5 distances at which these were presented, and 30 T only trials. Trial type was randomized within blocks, and inter-trial interval was random between 1250-2250ms (uniform distribution). Participants completed 10 to 12 blocks (∼10 minutes per block and ∼ 2h30 hours of total experimental time for a grand total of 3600 to 4320 trials; ∼2800 VT trials, 500 V trials, and 300 T trials) according to time constraints and were allowed to take brief breaks between blocks.

#### Behavioral Analysis

Behavioral analyses mimicked that described in Experiment 1, with the following exceptions. First, 7 distances were utilized as opposed to 6, and thus the Distance factor counts with an extra level. Further, due to the EEG recordings and the chosen approach to contrast evoked responses (i.e., with rapid on-off transients) (vs. induced; e.g., looming visual stimuli are turned-on rapidly but then sustained for a long duration), static and transient stimuli (vs. dynamic and continuous) were used in this experiment. In turn, tactile-only conditions could not be mapped onto particular visuotactile distances, and hence the initial ANOVA performed in Experiment 1 (contrasting VT and T trials as a function of distance) was not possible. Instead, we averaged VT RTs across all disparities and performed a paired-samples t-test between VT and T trials. Note that this approach is extremely conservative statistically, as in principle there should be no multisensory facilitation when visual and tactile stimuli are presented far from each other (e.g., Noel et al., 2015a,b; Salomon et al., 2017).

#### EEG Recording and Preprocessing

Continuous EEG was recorded from 128 electrodes with a sampling rate of 1000Hz (Net Amps 400 amplifier, Hydrocel GSN 128 EEG cap, EGI systems Inc.) and referenced to the vertex (Cz). Electrode impedances were maintained below 40 kΩ throughout the recording procedure. Data were acquired with NetStation 5.1.2 and further pre-processed using MATLAB and EEGLAB (Delorme and Makeig, 2004). Continuous EEG data were notch filtered at 60 Hz and bandpass filtered from 0.1 Hz to 40 Hz using an 8th order bi-directional zero-phase infinite impulse response (IIR) filter. Epochs from 200 ms before to 800 ms after stimuli onset were extracted and split according to experimental condition. Artifact contaminated trials and bad channels were identified and removed through a combination of automated rejection of trials in which any channel exceeded ±100 mV and rigorous visual inspection (e.g., Simon et al., 2017; Noel et al., 2019). A mean of 334.9 (SEM = 9.8) or 83.7% of trials were retained per VT condition, while 2.73 % (S.E.M = 1.71%) of channels were removed per participant. Data were then recalculated to the average reference and bad channels were reconstructed using spherical spline interpolation (Perrin et al., 1987). Lastly, data were baseline corrected for the pre-stimuli period (−200 to 0 ms post-stimuli onset).

#### EEG Analyses

Within the current study we adopt the so-called electrical neuroimaging framework (Brunet et al., 2011) for EEG analyses. Within this framework we leverage the fact that EEG recordings are performed from a full-montage of electrodes covering the entire skull and utilize data-reduction techniques to overcome the inherent multiple comparisons problem in EEG studies. Further, we avoid indexing particular components, which are both reference-dependent and potentially subject to experimenter bias. In turn, the global electric field strength present throughout the recording montage was quantified using global field power (GFP; Lehman & Skrandies, 1980; Lehman, 1987). This measure corresponds to the standard deviation of the trial-averaged voltage values across the entire electrode montage at a given time point, and represents a reference- and topographic-independent measure of evoked potential magnitude (Murray et al., 2008). Further, GFP is used as a data-reduction technique by summarizing 128 distinct time-series (i.e., electrodes) into a singular one.

In a first pass, we calculated average GFPs for each subject, as well as for the entire sample of participants (i.e., grand average) for tactile, visual, and visuotactile conditions separately while collapsing across distances. Time-resolved t-tests against zero were performed at each time-point from 200 ms pre-stimuli presentation to 800 post-stimuli onset in order to ascertain whether reliable evoked potentials were generated (to V, T, and VT stimuli). To account for the inherent auto-correlation problem in EEG, we set alpha at < 0.01 for at least 10 consecutive time points (Guthrie & Buchwald, 1991; see Noel et al., 2018c, d, and Simon et al., 2017, for a similar approach. Note that given the emphasis on GFP, a single time-course, we do not have a multiple comparisons problem requiring permutation testing).

Next, to ascertain whether a veritable multisensory effect existed (i.e., non-linearity between the co-presentation of V and T information vis-à-vis their presentation in isolation) we created visuo-tactile summed responses (hereafter, “summed” or “sum”) by adding the subject-level average responses to V and T. GFP was then calculated for this summed response and contrasted to the GFP of the multisensory visuotactile condition (or “paired” response; see Cappe et al., 2010 and Noel et al., 2018c for a similar approach). Indeed, as GFP is by definition positive, an advantage of utilizing this method within a multisensory framework is that supra- and sub-additivity indices may be measured (Sperdin et al., 2010; Murray & Wallace, 2012). The contrast between multisensory visuo-tactile discrepancies is undertaken solely for distances D2-D6, as the main interest here is in describing the neural correlates of PPS rapid recalibration and by definition there are no smaller visuo-tactile discrepancies than D1, and no larger discrepancies that D7. Having first established that the co-presentation of visual and tactile information resulted in a multisensory effect, we queried via a time-resolved one-way ANOVA whether VT GFPs differed as a function of visuo-tactile distance.

Lastly, having identified a time-period of interest (demonstrating both multisensory supra-additivity and space-dependent modulation of its multisensory response, see below) we examined whether this metric of PPS was altered as a function of the nature of the previous trial (i.e., trial t-1 being larger or smaller than t) and highlight the electrodes that are contributing to this effect.

### Experiment 3 - Neural Network Modeling

To suggest a putative mechanistic underpinning the observed rapid recalibration of PPS we employed a non-spiking biologically inspired neural network model that has previously been demonstrated to account for a number of PPS phenomena (Magosso et al., 2010a, b; Serino et al., 2015a; Noel et al., 2018b). Importantly, we did not attempt to build a new model from scratch to explain the rapid recalibration of PPS; contrarily, we simply took the most recent version of the model (Noel et al., 2018b) and imbued this model with Hebbian learning (as in Magosso et al., 2010a; Serino et al., 2015a) given the conceptual hypothesis that this form of learning ought in principle to account for rapid recalibration. This approach was taken as we considered it more powerful (conceptually) to demonstrate that a model already shown to account for a number of PPS phenomena can also incorporate the newly described rapid recalibration effect, than it is important to exactly fit behavioral results. Previous iterations of this model can account for sigmoidal facilitation functions (Serino et al., 2015a), the fact that PPS has different sizes for different body parts (Noel et al., 2018b), as well as its enlargement after tool-use (Magosso et al., 2010) and as a function of increasing exteroceptive signal velocities (Noel et al., 2018b). Thus, the model simply inherited previous parameters (see Table 1 in Noel et al., 2018a), with exception of those ruling Hebbian learning (for more detail regarding the model parameters and the robustness of the its PPS encoding to parameter selection see Noel et al., 2018b. For detail regarding Hebbian learning within PPS, see Serino et al., 2015a). In turn, here we only briefly explain the neural network implementation (for detail see Magosso et al., 2010a, b), and only briefly explore results from the simulation – focusing on the conceptual contribution, rather than the peculiarities of the model and its parameters.

The neural network simulates the peri-face space, although it could equally simulate PPS around any other body part (see Magosso et al., 2010a, Noel et al., 2018b). It includes two areas of unisensory neurons (tactile and visual) and a third area composed of a multisensory visuo-tactile neuron (see Figure 6A). Both the tactile and visual stimuli are mimicked by bidimensional Gaussian functions with small standard deviations (i.e. high precision in space) to simulate localized stimuli. Dynamic visual stimuli are simulated in the model thus resembling experimental conditions as in Experiment 1. Specifically, the visual stimuli are iteratively displaced closer and closer to the location of tactile receptive fields, to engender an approaching stimulus with equivalent velocity as that used in Experiment 1 (i.e., 75 cm/s for ∼200cm). Each unisensory area is composed by a matrix of N × N (N = 41) unisensory neurons. Unisensory neurons have a receptive field with a bidimensional Gaussian shape, through which the approaching stimulus is convolved (i.e., filtered), and are topologically aligned (i.e., proximal neurons respond to proximal spatial stimuli). The tactile unisensory neurons respond to tactile stimuli on the body part. Their receptive fields are 0.5 cm apart from one another along each dimension of the face, thus mapping a surface of 20 cm × 20 cm (approximate size of the face). The visual neurons respond to visual inputs in a planar space that extends sideways and in front of the body. Visual neurons’ receptive fields are 10 cm from one another along each dimension, mapping a space of 400 cm × 400 cm. Further, unisensory neurons within each area are reciprocally connected via lateral synapses (L) having a Mexican-hat pattern (near excitation and far inhibition).

Neurons within the two unisensory areas send excitatory feedforward synapses (*W*) to the downstream visuotactile area. This area mimics multisensory regions in the fronto-parietal cortex (e.g. ventral premotor cortex, ventral intraparietal area, area 7b), devoted to the representation of body-part specific PPS (e.g. peri-face). As electrophysiological data stress the existence of multisensory neurons having large receptive fields covering an entire body part, for parsimony only one multisensory neuron is included (see Magosso et al. 2010a, b; Serino et al. 2015a; Noel et al., 2018b, for a similar approach). The feedforward synapses from the tactile neurons to the multisensory one have a uniform value 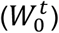. As such, the multisensory neuron has a tactile receptive field covering the whole face. The strength of the feedforward synapses from the visual neurons depends on the distance of the visual neurons’ receptive field from the body part. These synapses assume a maximum value 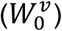 for visual neurons coding for the space covering and bordering the body part, then their value decreases exponentially as the distance of the visual neurons’ receptive field from the face increases. Finally, the multisensory neuron sends excitatory feedback synapses (*B*) to the tactile and visual unisensory neurons; the feedback synapses have the same pattern as the feedforward ones (see Noel et al., 2018b, Eqs 5-7 for more detail). Importantly, in the current work the strength of the feedforward excitatory connections from the unisensory neurons to the multisensory one are not set in stone, but are modified according to the following Hebbian rule:

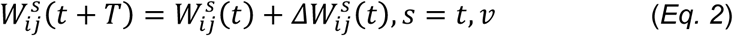

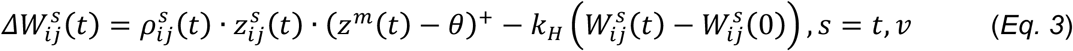

In Eq. 2 and Eq. 3, *ij* denotes the topographical position of a generic pre-synaptic unisensory neuron (t = tactile, v = visual) within the corresponding map, 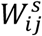 is the strength of the feedforward synapse connecting the presynaptic unisensory neuron at position ij with the post-synaptic multisensory neuron, 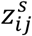 denotes the neural activity of the pre-synaptic unisensory neuron, *z*^*m*^ denotes the neural activity of the post-synaptic multisensory neuron, and (·)^+^ indicates the positive part of the function. *T* in Eq. 2 is the temporal step of synapses updating (*T* = 1 ms in our simulations), and *t* simple refers to a particular moment in the interval of the simulation. The Hebbian rule contains a reinforcing component (i.e., the first term in the right-hand member of Eq. 3) and a forgetting factor (i.e., the second term in the right-hand member of Eq. 3). According to the reinforcing factor, the strength of the feedforward synapse 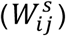 increases when both the pre-synaptic (unisensory) neuron and the post-synaptic (multisensory) neuron are active. In particular, the post-synaptic multisensory activity is compared with a small threshold *θ* (5% of the maximum activation), in order to avoid reinforcement in case of very small activity of the post-synaptic neuron. 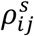 denotes the reinforcement learning factor, which is time-dependent (see below). The forgetting factor is constantly acting on the synapses regardless of neural activity, and is effectively inducing an exponential decay of the synaptic weight towards the fixed “basal” level 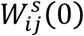 with a time constant 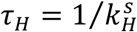 (in simulation steps). This time constant is a key parameter in the simulations as it determines the approximate timescale at which the network “remembers” past multi-sensory events. To avoid that the strength of excitatory connections increases unlimitedly, we imposed a saturation constraint for synapsis value; the reinforcement learning factor 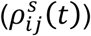 progressively reduces to zero as synapses approach their maximum value. We have:

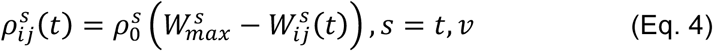

where 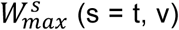 is the maximum value allowed for the tactile and visual synapses, and is assumed equal to the pre-existing (basal) value of the synapses on the face, that is 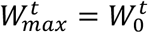 and 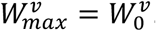. Therefore, tactile feedforward synapses (as well as visual feedforward synapses on and close to the face) are not subject to modifications. Value of parameter 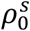 was assigned so that synapses reinforce gradually during stimulus presentation (i.e. several stimulation trials are required so that the reinforcing factor in Eq. 3 leads synapses close to saturation). The key parameter 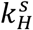 has been subjected to a sensitivity analysis to test how its value may affect a rapid trial-by-trial recalibration (see Section Results; see Noel et al., 2018b for sensitivity analyses of other parameters).

As in our previous models (Magosso et al., 2010a; Serino et al., 2015b), only feedforward synapses were subjected to training, while lateral synapses within unisensory areas and feedback synapses are kept fixed. This computational choice was made as in principle the training of the lateral and feedback synapses ought not to significantly affect our results (that would still mainly depend on feedforward synapses training). Feedback synapses can only be modified very seldomly, since their potentiation can easily produce phantom effects (i.e., activation in one unisensory area following stimulation in the other unisensory modality). The training of a large multitude of lateral synapses, on the other hand, requires an extremely delicate balance between overall excitation and inhibition to prevent network instability that can frequently occur during training. Most importantly, including a training of lateral synapses that maintains network stability (and spatial sensitivity) would result in a slight modification of the single bubble of activation in the unisensory areas, inducing yet even smaller effects on multimodal neuron activation and thus on PPS recalibration. In turn, in line with our previous studies (Magosso et al. 2010a; Serino et al. 2015b), we avoided to include these further mechanisms.

The overall input (say *u*) to a generic neuron (unisensory *s* = *t, v* or multisensory *s* = *m*) in the network is processed via a first-order temporal dynamics (Eq. 5, mimicking the post-synaptic membrane time constant) and a sigmoidal function (Eq. 6, mimicking the neuron’s activation function), generating the neuron’s output activity (say *z* (*t*)):

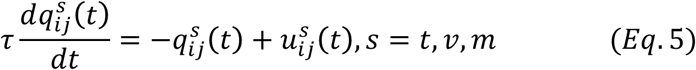

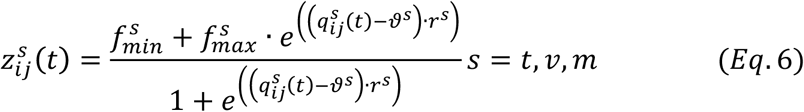

Eq. 5 and Eq. 6 hold for both unisensory *s* = *t, v* and multisensory *s* = *m* neuron; in case of the multisensory neuron (*s* = *m*) the equations hold without the subscripts as a single multisensory neuron is used in the network. Eq .5 describes the first order dynamics, where *q*^*s*^ (*t*) is the state variable, *u*^*s*^ (*t*) is the input to the neuron and*τ* the time constant. Eq .6 describes the sigmoidal activation function; 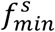 and 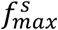 represent the lower and upper saturation of the sigmoidal function, *ϑ*^*s*^ establishes the central value of the sigmoidal function (i.e. the input value at which the output is midway between 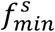 and 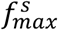) and *r*^*s*^ defines the slope. In turn, the output *z*^*s*^ (*t*) of each neuron is a continuous variable representing the particular neuron’s firing rate.

The overall input to the unisensory neurons (*u*^*s*^(*t*) s = t, v) is made up of the external input coming from outside the network (i.e., the stimulus filtered by the neurons’ receptive field *e*^*s*^(*t*) s = t, v), plus the lateral input coming from other neurons in the same area (via weights defined by lateral synapses *L*), and feedback input from the multisensory neuron (via weight defined by the feedback synapses *B*, see Eq. 7). The overall input to the multisensory neuron is made up of the feedforward inputs from the two unisensory areas (via weights defined by the feedforward synapses *W*, see Eq. 8).

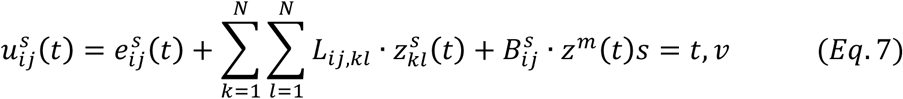

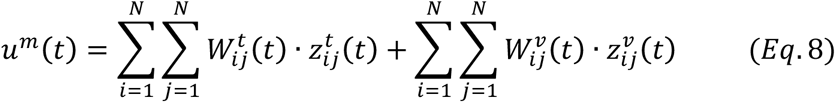

where *L*_*ij,kl*_ in Eq. 7 denotes the strength of the lateral synapse from the pre-synaptic neuron at position *kl* to the post-synaptic neuron at position *ij* within the same unisensory (tactile t or visual v) area, and 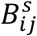 is the strength of the feedback synapse from the multisensory neuron to the unisensory neuron at position *ij* within the tactile (t) or visual (v) area. The sums in Eq 7 and 8 extend to all neurons within each unisensory area.

Lastly, since behavioral data are expressed in terms of tactile reaction times, we decoded such a measure from the network. This network tactile RT was computed as the time necessary for the overall tactile activity (the sum of all tactile neurons’ activity) to reach a given threshold *P*_*th*_= 4 starting from the tactile stimulus onset. Since neuron activity ranges between 0 and 1, this means that an ensemble of a few tactile neurons needed to be active for the stimulus to be detected. In the multisensory condition, the activation in the tactile area can be speeded up compared to the unisensory condition, and thus the network RT decreased, when the visual stimulus is able to trigger the multisensory neuron. For each condition (unisensory, and multisensory at each of the visuo-tactile disparities and), twenty trials were simulated.

## Results

### Augmented Reality Psychophysics – Experiment 1

Overall participants were very accurate at withholding responses during unisensory visual catch trials (< 1% false positives). The initial 2 (Condition: Experimental vs. Baseline) X 6 (Distance: D1 through D6) within-subjects ANOVA demonstrated a significant main effect of Condition (F(1, 37) = 10.92, p = 0.003, η2 = 0.313; Experimental, M = 0.289 s, S.E.M = 0.010 s; Baseline, M = 0.333 s, S.E.M = 0.011 s) and a significant main effect of Distance (F(5, 190) = 2.324, p = 0.047, η2 = 0.088). Importantly, and as illustrated in Figure 1B, results revealed a significant interaction between these factors (F(5, 190) = 3.463, p = 0.005). This interaction is further explained by the lack of a main effect of Distance in the Control condition (F(5, 190) = 0.668, p = 0.649), and the presence of this same effect in the Experimental condition (F(5, 190) = 46.97, p < 0.001, η2 = 0.559). Thus, participant’s RT to tactile stimulation became faster the closer a task-irrelevant visual stimulus was to their body, and this result cannot be explained merely by an expectancy effect (see Kandula et al., 2017). The central point of the sigmoidal function best describing multisensory RTs as a function of visuo-tactile distance was at D = 3.74 (see Figure 1B).

With regard the fitting procedure, when trials were divided given the nature of their precedent trial, goodness of fit measures demonstrated that for both conditions (T-1 Smaller and T-1 Larger) the sigmoidal fitting described the data equally well (R^2^ T-1 Smaller = 0.84, R^2^ T-1 Larger = 0.81, paired-samples t-test p = 0.62). The central point, describing the spatial extension of PPS representation, was significantly modulated as a consequence of the nature of the immediately preceding trial, as demonstrated by a significant difference between T-1 Smaller (M = 3.42, S.E.M = 0.11) and T-1 Larger (M = 4.04, S.E.M = 0.13; t(27) = 3.6707, p < 0.001; Figure 1C). We did not statistically compare the central point of T-1 Smaller and T-1 Larger with that of the central point when trials were not split depending on sensory history, as this comparison would be confounded by a significantly different amount of repetition in each condition. However, it must be noted that numerically, the extension of PPS when not split as a conditional of T-1 falls in between the values reported for T-1 Smaller and T-1 Larger. Although there was a trend, statistically there was no significant difference in the value of the slope (p = 0.17) describing visuo-tactile RTs as a function of whether the precedent trial had been larger or smaller. Lastly, given that a number of participants were excluded from this analysis given poor fits in one of the conditions, we confirmed the above-mentioned results via a paired t-test contrasting unisensory and multisensory responses, as well as a 2 (T-1 Smaller vs. T-1 Larger) x 4 (Distance; D2-D5) within-subjects ANOVA including all 38 subjects. These latter analyses confirmed that multisensory reaction times were faster than unisensory ones (t = 6.71, p = 7.03×10^−11^), that visual proximity facilitated tactile reaction times (F = 5.04, p = 0.02), that generally reaction times after T-1 Larger were quicker (F = 7.69, p = 0.009) than after T-1 smaller trial, and most importantly that trial-to-trial sensory history remapped PPS (F = 3.47, p = 0.04; see Figure S1). Analysis focused solely on the 17 participants removed from the sigmoidal fit analysis also showed a multisensory vs. unisensory main effect (t = 2.82, p = 0.03), a facilitation of tactile reaction times when visual stimuli were near (F = 2.88, p = 0.04), and a significant interaction between these variables (F = 2.03, p = 0.03).

Given the rapid recalibration effect, we defined the degree at which each participant rapidly recalibrated his/her representation of PPS as the difference between their central point value for T-1 Larger and T-1 Smaller trials (PPS Rapid Recalibration = T-1 Larger – T-1 Smaller), and then correlated this value with the particular participant’s *b* value (that contributes to define the raw PPS slope, see Eq. 1). This analysis was motivated by the audio-visual rapid recalibration literature (Van der Burg et al., 2013; Noel et al., 2016a) which demonstrates a strong relation between the amount a particular participant rapidly incorporates sensory history and their a priori sensitivity to the task at hand. Similar to the audio-visual studies, our analyses (R^2^ = 0.22, p = 0.03; Figure 1D) indicated that the shallower, or more gradual, a participant’s gradient between ‘near’ and ‘far’ space representation (i.e., the less well defined a participant’s PPS boundary is, corresponding to flatter slopes), the *more* he/she will recalibrate his/her PPS representation. Please note that the relationship is positive here, and not negative, as in the sigmoidal function a larger *b* value (see Eq.1) indicates a shallower slope (parameter *b* contributing to defining the steepness/shallowness of the sigmoidal function).

### Psychophysics during EEG – Experiment 2

As for Experiment 1, overall participants were very accurate at withholding responses during catch trials (false alarm on 3.75% of trials, S.E.M = 1.20%), and thus behavioral results are analyzed solely in light of RTs (see Noel et al., 2015a, 2015b). The paired-samples t-test between visuotactile multisensory presentation and tactile alone presentations was significant (t = 6.76, p = 1.4e-06), indicating that responses to the former condition were quicker (M = 296ms, S.E.M = 9.8ms) than to the latter (M = 317ms, S.E.M = 7.3ms). Namely, the co-presentation of visual stimuli with tactile stimulation resulted in multisensory facilitation.

Next, we fit visuo-tactile RTs to a sigmoidal function (see Methods and Experiment 1) on an individual subject level (goodness of fit; r^2^ = 0.76) and extract the central point and slope of the function at the central point. This procedure indicated that on average the inflection point where visual stimuli facilitated tactile RTs was D = 1.73 (see Figure 2B). Having divided trials given the nature of the immediately precedent trial (i.e., whether a smaller or larger visuo-tactile disparity has been presented on trial T-1) we contrast both the central point and slope of the function describing tactile RTs given visuo-tactile distance as a function of trial history via a paired samples t-test. This analysis demonstrated that when the precedent trial had been one in which a smaller visuo-tactile spatial disparity was indexed, PPS was smaller than when on the previous trial a larger visuo-tactile disparity had been probed (central point when T-1 smaller, M = 2.83, S.E.M = 0.11; central point when T-1 larger, M = 4.42, S.E.M = 0.20; t-test T-1 smaller vs. larger, t = 7.76, p = 1.83e-7; Figure 2C). Thus, the same effect found as in Experiment 1 for face PPS was replicated behaviorally in Experiment 2 for hand PPS. There was no difference in the *b* parameter (Eq. 1, contributing to slope), although again a trend existed (p = 0.07) for the *b* value becoming larger after a large visuo-tactile disparity (T-1 larger, *b* = 9.19 +/-1.97, T-1 smaller, *b* = 4.57 +/-1.59). Lastly, given the results from Experiment 1 we correlated the amount a particular subject’s PPS shifted due to rapid recalibration (i.e., central point when T-1 larger minus central point when T - 1 smaller) and the parameter contributing to defining the slope (*b*). Contrary to Experiment 1 in this case we did not find a linear relationship between these variables (r = −0.22, p = 0.32).

### Electroencephalography Results

#### Multisensory Responses

Time-resolved t-tests of GFP values to zero (i.e., baseline) indicated significant evoked responses to T (69-93ms post-stimuli onset and 167ms post-stimuli onset and onward), V (75-89ms post-stimuli onset as well as 96ms post-stimuli onset and onward), and VT (78ms post-stimuli onset and onward) stimuli (see Figure 3 left column). Hence, in a subsequent step we directly contrasted the paired multisensory response with an artificially created summed visuo-tactile condition that counted with the same energies presented (e.g., V+T), but were not concurrently presented (Figure 3, right column). This analysis revealed two transient time-periods demonstrating supra-additivity (i.e., VT>V+T; t1, 124-158ms post-stimuli onset; t2, 204-223ms post-stimuli onset), as well as a more sustained epoch of sub-additivity (i.e., VT<V+T; 315ms post-stimuli onset onward). As an additional analysis, to control for the fact that the experimental design included a greater number of VT than V or T trials – leading to putative differences in signal-to-noise ratios – we subsampled individual subject data to match trial numbers and repeated this analysis. The results were virtually identical, showing supra-additivity from 123-160ms and 204-226ms post-stimuli onset (see Supplementary Material, Figure S2). The different number of VT, V, and T, trials could have also led to the less frequent unisensory stimuli being processed as oddballs (Squires et al., 1975), yet PPS processing has been shown to be independent of attention (Salomon et al., 2017), and somehow one would have to explain why the oddball condition is not modulated by distance, while the standard (VT) condition is.

**Figure 3.**
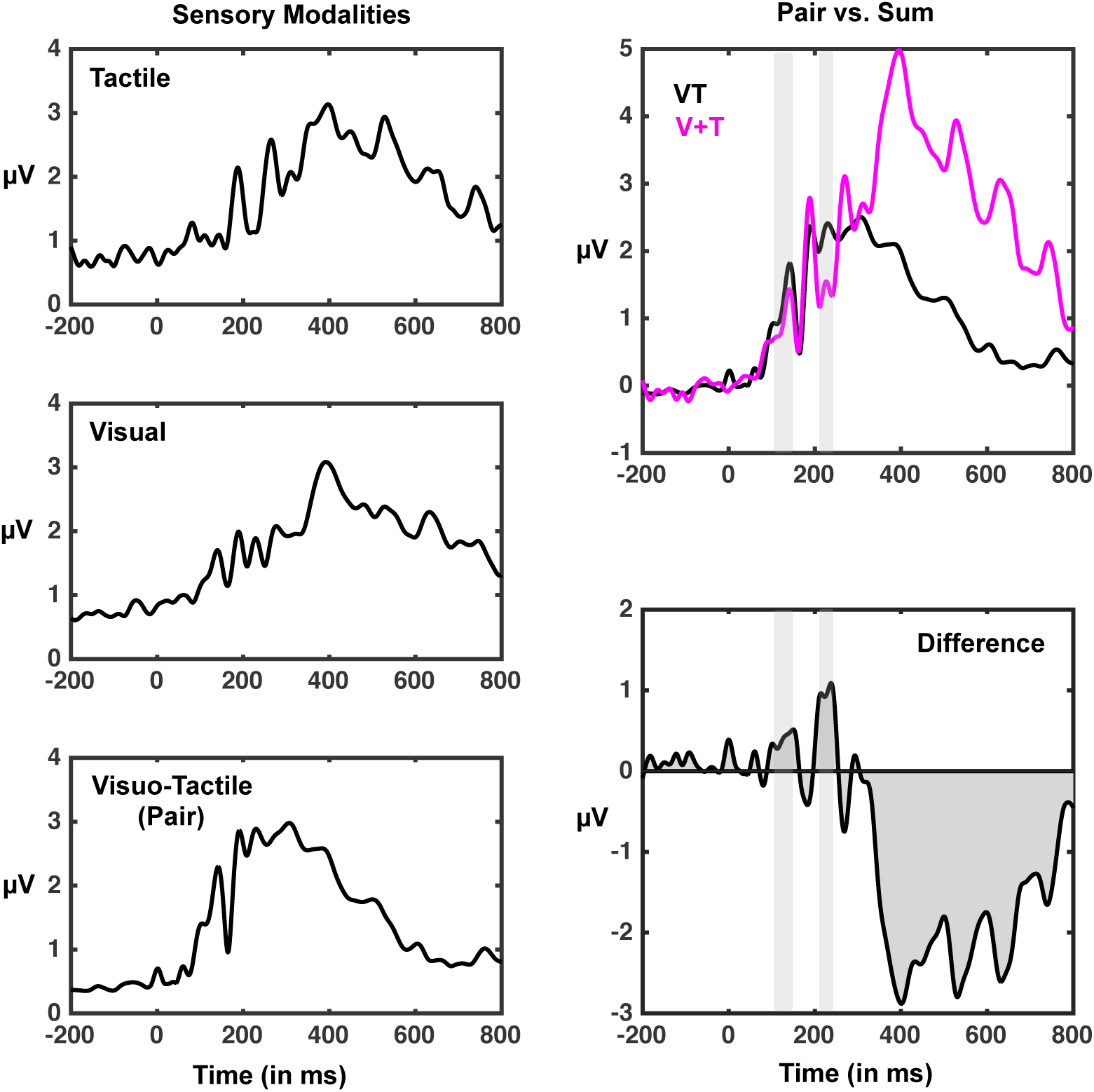
Global Field Power associated with distinct sensory stimulations. Left-panel; GFPs associated with tactile (top), visual (center), and visuo-tactile (bottom) stimulation. Right-panel; contrast between paired (VT; black) and summed (V+T; purple) GFPs (y-axis) as a function of time since stimuli onset (x-axis). Top panel demonstrated the raw values associated with both paired and summed conditions, while the bottom panel is the difference wave (pair – sum) between the two. Thus, positive values indicate supra-additivity, while negative values indicate sub-additivity. Gray shaded areas correspond to time-periods where supra-additivity is significant (p<0.01). See Figure S2 for a similar analysis while subsampling trials to match across conditions.

#### Space-Dependent Multisensory Responses

Given that GFP analyses contrasting multisensory and unisensory responses revealed instances of true multisensory integration (i.e., non-linearity), we next examined whether multisensory responses were modulated by visuo-tactile distance. A one-way ANOVA contrasting visuo-tactile distances D2, D3, D4, D5, and D6 showed a significant effect of distance between 70-191ms post-stimuli onset. Interestingly, a comparison of GFPs averages across the 130-150ms post-stimuli onset interval (an interval demonstrating the significant one-way ANOVA effect and exhibiting a clear peak in GFP) revealed a monotonic effect where GFP was largest for the nearest distance (D2, M = 2.52, SD = 0.30) and then in sequence D3 (M = 2.25, SD = 0.30), D4 (M = 2.10, SD = 0.28), D5 (M = 2.04, SD = 0.26), and D6 (M = 1.79, SD = 0.28). All pairwise comparison were significant from one another (p<0.001), except for the contrast between D4 and D5 (p = 0.068; Figure 4). Importantly, the same analysis contrasting differences for the visual condition alone did not show a significant main effect (F = 1.17, p = 0.32, see Figure S3). At a sensor and voltage level, the electrodes driving this GFP spatial effect were clustered over the occipital cortex and frontal cortex bilaterally (see Figure 4). This GFP landscape, which is largely driven by a posterior positivity (see Figure 5A), is consistent with a visual P1 (Di Russo et al., 2002). That is, seemingly the modulation as a function of distance is reliant on the presence of a tactile signal, but the major driver of EEG signals around 100-150ms post visuo-tactile stimulus onset is visual.

**Figure 4.**
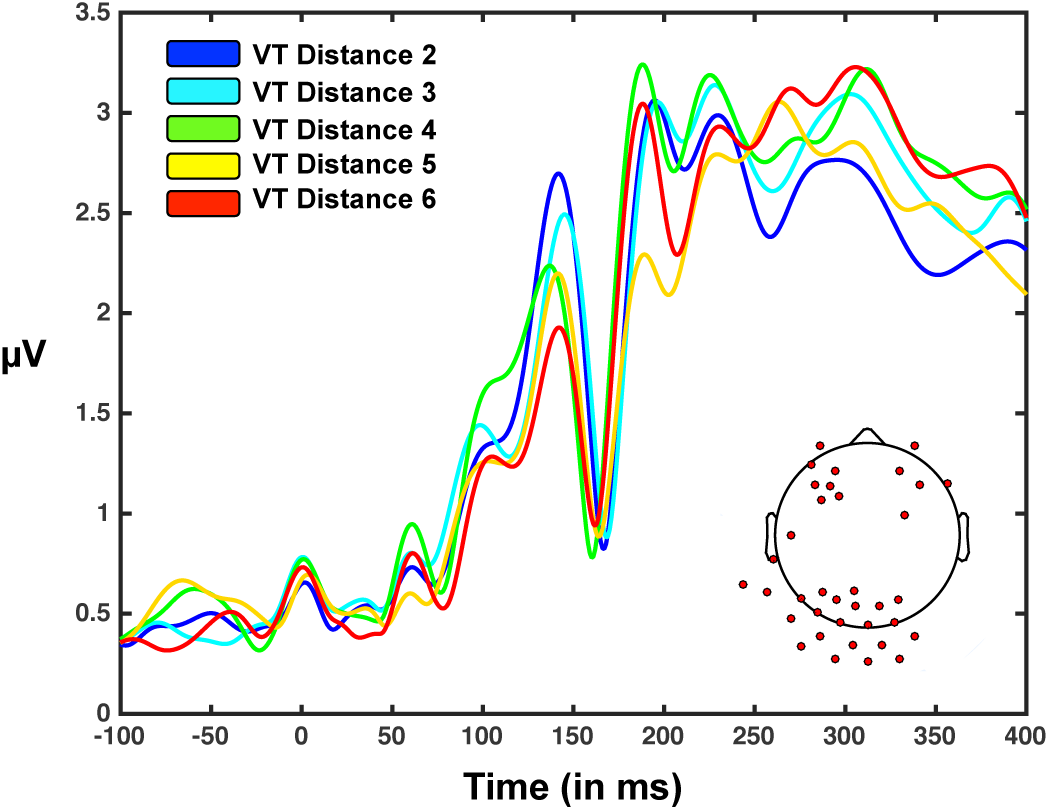
Visuotactile GFP as a function of visuo-tactile distance (D2 = nearest; D6 = farthest). GFP (y-axis) as a function of time from stimuli onset (x-axis) and visuo-tactile distance (from nearest to farthest; D2 = blue, D3 = cyan, D4 = green, D5 = yellow, D6 = red). Bottom right insert illustrated electrodes showing a one-way ANOVA distance effect at the voltage level, and hence driving the GFP difference. These electrodes cluster in the occipital cortex (one continuous cluster), as well as bilaterally in the frontal lobe (separate clusters). See Figure S3 for a similar analyses for the visual-only condition, which shows no modulation of the GFP peak emphasized here as a function of distance.

**Figure 5.**
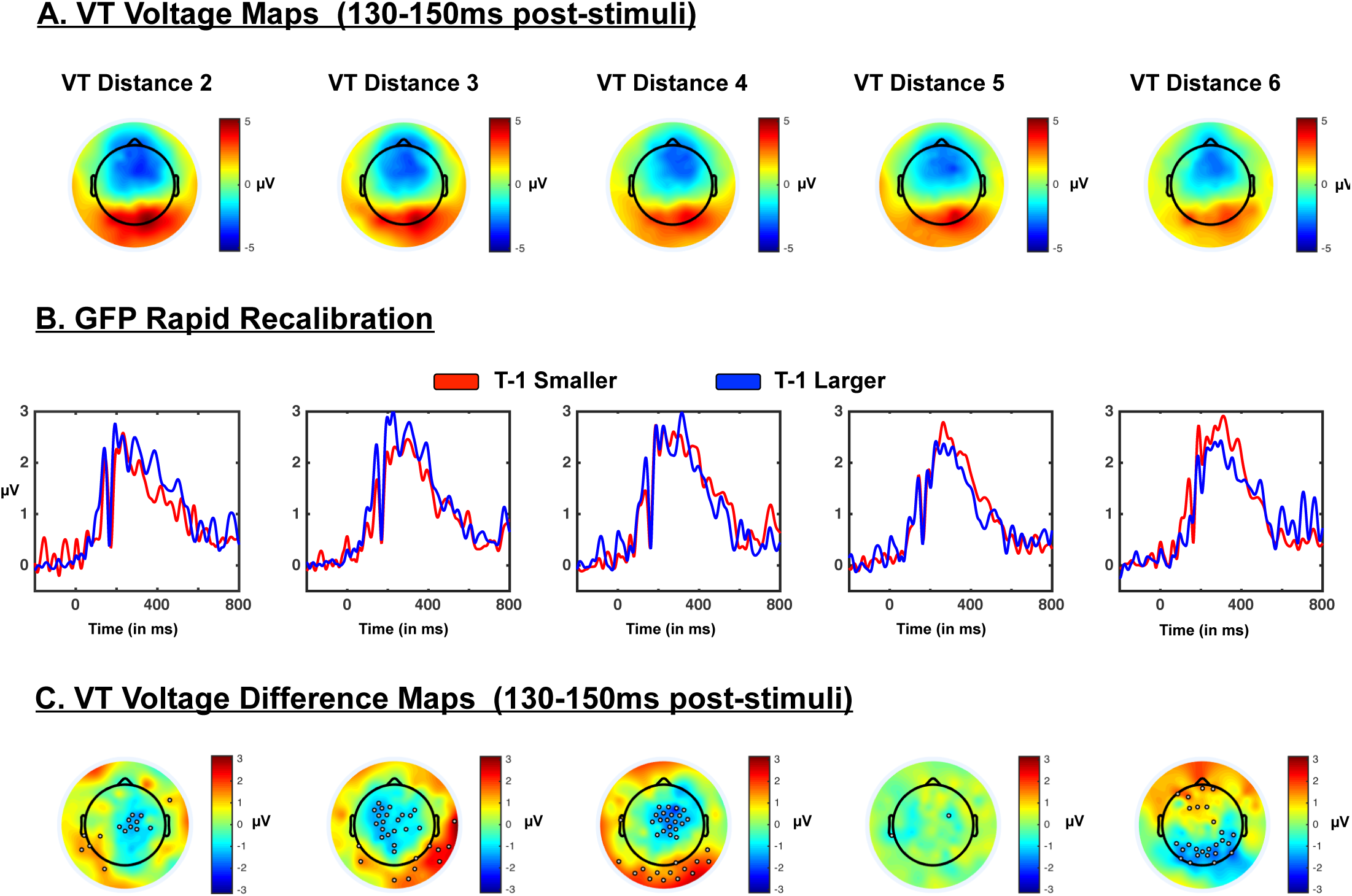
Neural correlates of PPS rapid recalibration. A) Topography of voltages during the time-period indicating multisensory supra-additivity and space-dependency in VT trials. Maps indicate that the major driver of neural activity between 130-150ms post-stimuli onset has a positive component at occipital electrodes and negative component in frontal sensors. Topographies are similar across distances (from D2 to D6). B) GFP as a function of distance and nature of the immediately precedent trial. When trial t-1 was larger (blue) than the currently indexed distance (e.g., D5 at t-1 and D3 at t) GFPs between 130-150ms post-stimuli onset are seemingly larger at distances D3 and D4, than when the previous trial was one with a smaller (red) visuo-tactile disparity. C) Topography of voltage difference between t-1 smaller and t-1 larger during the time-period indicating multisensory supra-additivity and space-dependency in VT trials. The topographical distribution of the difference in GFP when t-1 was smaller vs. larger than the current visuo-tactile discrepancy seemingly indicates that rapid recalibration of PPS is driven by centro-occipital sensors.

**Figure 6.**
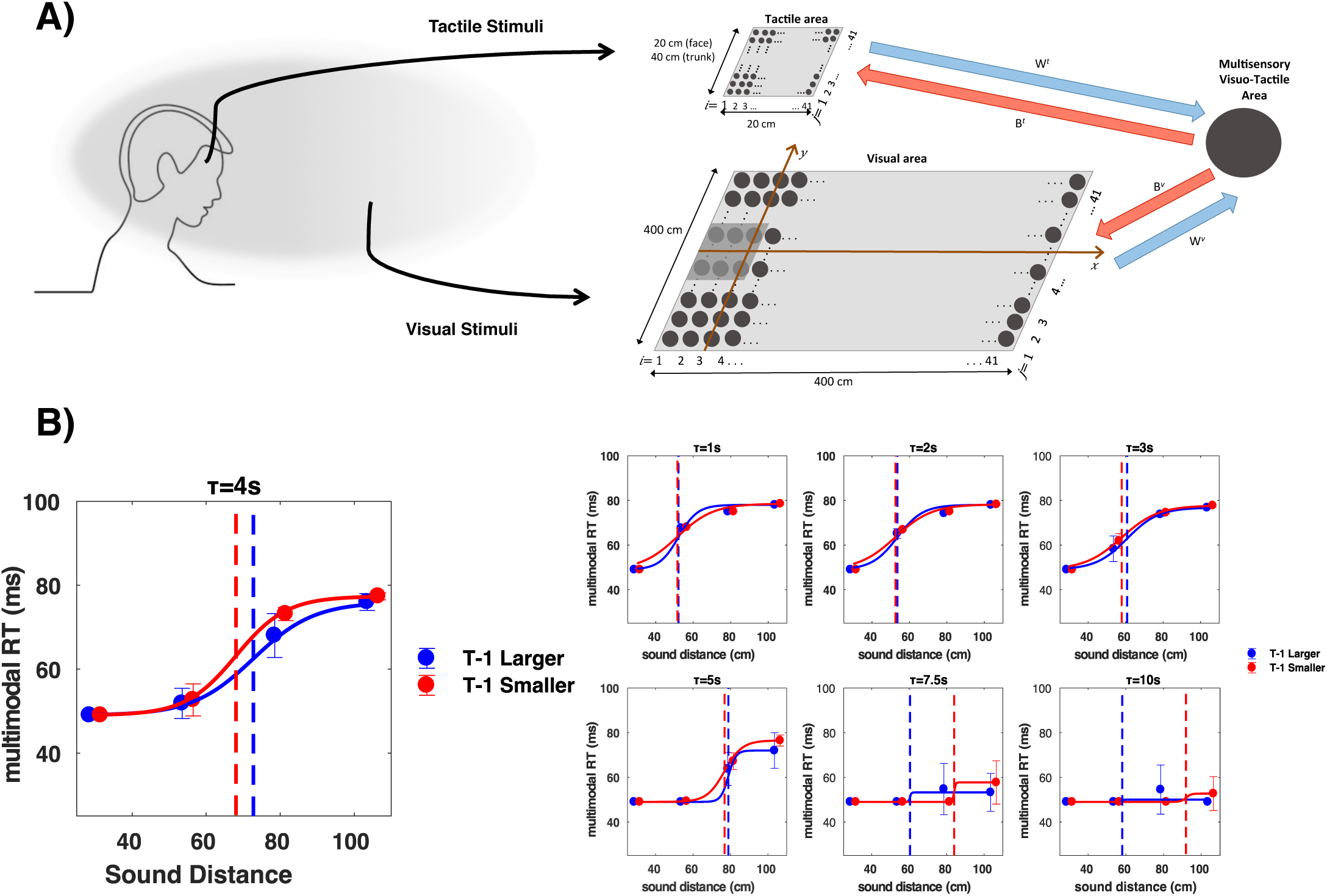
Hebbian learning within a neural network of PPS can account for rapid recalibration. **A)** Schematic of the model; tactile neurons have receptive fields encoding for the face, visual neurons encode for the external space. Neurons within areas are connected to one another via a Mexican hat pattern, and project to a multisensory neuron. Visual neurons closer to the location of touch have stronger projections (see text). The multisensory area projects back to unisensory areas, and connections are not set in stone, but strengthen and weaken according to Hebbian learning – whether pre- and post-synaptic neurons fire together or not. **B)** Results simulating reaction times to touch as a function of visuo-tactile proximity given the neural network model, Hebbian learning, and sensory history.

#### Rapid Recalibration of Space-Dependent Multisensory Responses

Having established that supra-additive multisensory effects were present within the interval ∼130-150 ms post-stimuli onset, and that this interval equally demonstrated a space dependency, we next examined whether we could discern a neural correlate to the rapid recalibration effect. Figure 5A illustrates the topography of voltages across the entire montage during the time-period of interest, suggesting a visually driven dipole (positivity in occipital sensors will lead to a negative pole in frontal sensors given a full montage). Next, the data were categorized as a function of whether the previous trial had been one in which a larger or smaller visuo-tactile discrepancy than the current was presented. A 2 (T-1 smaller vs. T-1 larger) x 5 (distance; D2-D6) ANOVA on the mean evoked GFP present between 130-150ms post-stimuli onset in the VT condition demonstrated main effects of distance (F(4, 104) = 151.56, p < 0.001) and the nature of previous trials (F(1, 26) = 70.42, p < 0.001), as well as an interaction between these variables (F(4, 104) = 117.73, p < 0.001). As indicated above the main effect of distance was due to a monotonic reduction in GFP as a function of distance, while the main effect of trial history was due to larger GFPs when the precedent trial had been a large spatial disparity (M = 2.22, S.E.M = 0.051) than when the precedent trial had been one with a smaller spatial disparity (M = 2.05, S.E.M = 0.053). Most importantly, the interaction was driven by significant difference between T-1 smaller vs. T-1 larger at D3, D4, and D6 (all p < 0.001), but the lack thereof at D2 and D5 (all p > 0.10). The difference at D3 (T-1 larger, M = 2.60, S.E.M = 0.35; T-1 smaller, M = 1.90, S.E.M = 0.29) and D4 (T-1 larger, M = 2.48, S.E.M = 0.30; T-1 smaller, M = 1.82, S.E.M = 0.32) were due to stronger evoked responses in the T-1 larger condition (vs. T-1 smaller), while the opposite was true at D6 (T-1 larger, M = 1.38, S.E.M = 0.26; T-1 smaller, M = 1.98, S.E.M = 0.30). Figure 5B illustrates the time-course of the rapid-recalibration effect. Importantly, Figure 5C highlights the electrodes driving the difference in GFP (difference between T-1 smaller vs. larger) within the time period between 130ms and 150ms post-stimuli interval. Interestingly, while the GFP is dominated during this time-period by a visual response (Figure 5A, P1, see above), the modulation of this response as a function of immediately precedent visuo-tactile disparity is seemingly majorly driven by central electrodes, putatively indexing a somatosensory response. The distinct contrast topography (T-1 Smaller – T-1 Larger) for D6 (vs. D3 and D4) appears to be driven by the lack of a consistent response when T-1 was larger on the previous trial (see blue curve in Figure 5B). At D6 there are few trials where the previous disparity was larger (solely D7), and this could contribute to the lack of robust response. The same is true for smaller disparities than D2 (solely D1), but at this distance the VT responses were generally stronger (given the main effect described above).

### Neural Network Model

In a naturalistic and ecologically valid augmented-reality setup we demonstrate that PPS recalibrates on a trial-by-trial basis, and then we replicate this behavioral effect with static as opposed to dynamic stimuli, while participants’ electroencephalogram is monitored. The neural results suggest that visuo-tactile responses are supra-additive, and these responses, but not visual-only, are graded as a function of distance during the time-period spanning 130 and 150ms post-stimulus onset. This suggests that the multisensory EEG response indexed is not related to absolute visual distance, but disparity between touch and vision. Further, the fact that tactile stimulation, and importantly its history, modulate the same “mostly-visual” response, suggests a local, perhaps microcircuit level, mechanism. That is, the short-term hysteresis demonstrated does not appear to depend on, say, a long-range feedback connection from higher-order areas – this would have been evidenced as T-1 Smaller vs. Larger differences occuring earlier than 130ms. In turn, we hypothesized that rapid recalibration of PPS may be driven by a change in the strength or pattern by which visual and tactile information converge.

In turn, we employed a well-established neural network model of PPS (see Figure 6A) and asked whether – in principle – Hebbian learning (Hebb, 1949) could account for the rapid recalibration of PPS. In particular, the strength of synapses was not set in stone, but was allowed to alter within the cadre of Hebbian learning, and given sensory stimuli mimicking those presented in Experiment 1 (i.e., looming stimuli at 75cm/s and with 2000 ms between the onset of each approach). In the model, the strength between different unisensory neurons and the multisensory node increased when both the pre-synaptic (unisensory) neuron and the post-synaptic (multisensory) neuron were concurrently active. Further, all synapses were subject to exponential forgetting (to avoid a scenario where all synapses are maximal; see *Methods* for further detail). As a first attempt to determine whether our inherited neural network model of PPS (Magosso et al., 2010a; Noel et al., 2018b) could account for the behavioral results presented here, we simulated a psychophysical experiment with the time constant of the network forgetting rate set to *τ*_*H*_ = 4*s*.This parameter was set based on the intuition that, in order for the recalibration of PPS to be effective, the time constant of the forgetting rate must be of the same order of magnitude as the time that separates two consecutive responses. Since the meaningful learning takes place at the moment of tactile stimulation, if the time constant of the network is much smaller than the time between stimuli, the network goes back to its basal state and synapses before the following trial and no recalibration can be observed. If the time constant is much larger, instead, the network will retain information from several preceding trials, therefore synapses will continue to reinforce until they are all saturated, effectively eliminating the PPS altogether (i.e., no space-dependent effect). As illustrated in Figure 6B, the model with *τ*_*H*_=4s could indeed replicate the above-mentioned psychophysical effects, with a difference in the central point of the sigmoid of 4.5 cm between T-1 larger and T-1 smaller trials. Interestingly, and similarly to what observed behaviorally, most of the difference is observed in the far space. In the simulations, this happens because when the stimulus is close, the multisensory neuron is already close to maximally active solely because of the visual stimulus. Therefore, the tuning of visual synapses that are close to the body cannot be further enhanced given sensory history. Behaviorally, this is reflected by a floor effect on reaction times, which are already maximally fast when the current stimulus is close, regardless of previous history.

To further test our hypothesis regarding the role of the forgetting rate – and provide an analysis regarding the sensitivity of the network to changes in this parameter - we performed further simulations in which we set *τ*_*H*_ to a set of values spanning between 1 and 10s. To be able to compare the results across values of *τ*_*H*_, the simulations were performed on the same set of permuted stimulation distances, as the order of trials influences the results. For the majority of *τ* values tested, 2 < *τ*_*H*_ < 7.5*s*, the differences we observe in the RT curves are in the expected direction (equal to that reported for *τ*_*H*_ = 4). Also in line with our predictions, there was little to no change as a function of sensory history when *τ*_*H*_ ≤ 2*s*, this is because the network forgets quickly. Lastly, when*τ*_*H*_ > 7.5*s* the model yields a flat PPS, that is, no real PPS. This last result suggests that the rapid recalibration of PPS is not solely an interesting oddity, but within this Hebbian framework may in fact be a crucial component to having a PPS in the first place: the time constant *τ*_*H*_ ruling the forgetting factor of synaptic plasticity must be within a defined range to allow for a i) a PPS, ii) that is sensitive to environmental factors. In other words, for the vast majority of temporal constants allowing for PPS (*τ*_*H*_ < 7.5*s*), rapid recalibration occurs (solely when *τ*_*H*_ < 2*s*, there is no rapid recalibration).

## Discussion

We noted that PPS is routinely argued to remap adaptively, given the state of the sensory environment. PPS has been shown to re-size or re-shape after training lasting a few minutes (Farne & Ladavas, 2000; Maravita et al., 2000; Canzoneri et al., 2013), a few hours (Bassolino et al., 2015), months (Iriki et al., 1996), or years (Serino et al., 2007). Importantly, however, the timelines of this recalibration – from a few minutes to years - does not match an adaptive time-scale. Here we questioned whether PPS remaps on a trial-by-trial basis as a function of short-term sensory input and history.

In Experiment 1 we measured in an ecologically valid augmented reality setup (Serino et al., 2017) the distance at which a dynamic external (visual) stimulus affected tactile processing of touch administered on the cheek. This space-dependent multisensory facilitation of tactile processing is taken to be an index of PPS (Serino, 2019). We then analyzed tactile responses based on the type of multisensory stimulation received on the immediately preceding trial, i.e., whether touch was given when the visual stimulus was either closer or farther from the participant with respect to the current trial. PPS was respectively larger or smaller when the preceding trial implied a multisensory interaction at a farther or at a closer distance, suggesting that PPS rapidly recalibrates based on prior sensory statistics. Experiment 2 successfully replicated the same rapid recalibration for the peri-hand space (in an independent set of participants and a different experimental setup), and demonstrated the generalizability of the findings from Experiment 1, by showing trial-to-trial recalibration while static (vs. dynamic) and physical (vs. virtual) visual stimuli were presented concurrently to tactile stimulation on the hand (vs. face). Further, while a considerable number of participants showed poor sigmoidal fits in Experiment 1 and thus were excluded from that analysis (although included in confirmatory analyses of variance), in Experiment 2 behavioral fits were good for all but one subject (likely due to the greater number of repetitions) and thus provides strong support for the replicability of the behavioral finding. Together, the psychophysical results argue that PPS may not only remap dynamically – that is, *within* a trial (Brozzoli et al., 2009, 2010; Patane et al., 2019) – or plastically on a slow time-scale (Iriki et al., 1996; Maravita & Iriki, 2004; Farne & Ladavas, 2000; Canzoneri et al., 2013) – but that a third mode exists; rapid plasticity.

Electroencephalographic monitoring of the visuo-tactile task (Experiment 2) showed that the Global Field Power (GFP) approximately 130ms post visuo-tactile presentation is supra-additive (i.e., demonstrates multisensory integration) and scales with visuo-tactile proximity (i.e., demonstrates a PPS effect). Interestingly, the amplitude of this same response (∼130ms post-stimuli onset) was altered as a function of whether the previous trial had been one with greater or smaller visuo-tactile disparity. The timing of this effect is in line with descriptions of the EEG components present during rapid recalibration of audio-visual temporal acuity (Simon et al., 2017). The voltage response at this time-period was seemingly visually driven, while the topography of its modulation (i.e., difference wave) suggested an additional central origin, likely sensorimotor (yet this is speculative given the inverse problem). The conjecture that the history dependent modulation in visual P1 amplitude was driven (at least in part) by sensorimotor signals is further supported by the fact that this modulation was most evident at distances showing a PPS remapping (from D2/D3 to D4/D5), as defined by tactile reaction times. The rapid recalibration topography also suggests potential voltage amplitude differences at occipital sensors, which would be in line with recent demonstration of bottom-up early multisensory integration in classically considered unisensory areas (see Ghazanfar & Schroeder, 2006 and Schroeder & Foxe, 2005 for reviews).

Overall, these results add to the general neuroimaging literature on PPS (see Grivaz et al., 2017 for review) and the burgeoning study of this multisensory space in a time-resolved manner. Sambo & Foster, 2009, showed modulations in the amplitude of the visual P1 and N140 with tactile proximity, but they only indexed two distances and did not ascertain the multisensory – integrative – nature of this response, nor if this response mimicked spatial remapping. In recent studies, Bernasconi and colleagues (2018), as well as Noel and colleagues (2019), showed a modulation of local field potentials evoked by touch on the trunk/hand as a function of the proximity of auditory stimuli presented at different distances. Results demonstrated PPS processing (i.e., different response for near vs. father stimuli) as early as 50ms post-stimuli onset, from insular cortex, but most commonly around 200ms post-stimuli onset, mainly from pre- and post-central gyri (Bernasconi et al., 2018; Noel et al., 2019). These studies, however, did not include a condition wherein PPS was remapped, and thus there has been no description of the EEG correlates of PPS remapping. Lastly, Noel and colleagues (2018b) showed that audio-visual responses are facilitated near the boundary of PPS, and only evident ∼300ms post-stimulus onset. Together, this pattern of results is evocative of a temporal cascade, wherein visuo-tactile near space is differentiated from the far space first (∼130ms), then audio-tactile space is (∼200ms), and finally perhaps a space-selective audio-visual processing (Van der Stoep et al., 2016; Noel et al., 2018d) is scaffolded upon this encoding of PPS. This speculation, however, needs to be confirmed by further studies directly comparing visuo-tactile, audio-tactile, and audio-visual integration as a function of distance.

The neuroimaging results hint at two interesting possibilities. First, the P1 is considered to index fidelity of visual encoding (Di Russo et al., 2002), and hence the fact that when paired with tactile stimuli this component is moderated by visuo-tactile distance implies that visual perception itself may be different within and outside the PPS. Indeed, initial studies in this domain have suggested that shape discrimination is better within the PPS – even after accounting for relative size (Blini et al., 2019). Second, the fact that the index of rapid recalibration of PPS was *not prior* to the component graded by visuo-tactile disparity (∼130ms) arguably suggests that this remapping is driven by a local re-weighting of the convergence between visual and tactile signals, or the co-activation of both somatosensory and visual responsive units. In this line, we have recently suggested that dynamic remapping, at least that provoked by stimuli velocity, may be instantiated by firing-dependent adaptation of multisensory neurons (Noel et al., 2018b). Speculatively, the mechanism behind the slower timescale plasticity of PPS most often indexed in the PPS literature may originate from distinct neural areas (e.g., feedback connections from arousal or valence centers during value-based remapping; Bufacchi & Iannetti, 2018). That is, we hypothesize that the faster timescale remapping of PPS (dynamically within a trial, or rapidly on a trial-by trial basis) may be intrinsic to the multisensory PPS circuitry (i.e., respectively, neural adaptation and Hebbian re-weighting of synapses), while slower and longer-lasting recalibration may be driven by factors extrinsic to the PPS circuitry itself (Bufacchi & Iannetti, 2018; see Noel & Serino, 2019).

To further ascertain whether Hebbian plasticity is in principle capable of accounting for rapid recalibration we employed a well-established neural network model of PPS (Magosso et al., 2010a, b). In particular, we added to the model presented in Noel et al., 2018b – which is capable of dynamic resizing due to neural adaptation – the flexibility of strengthening or weakening the feedforward synapses, i.e. the synapses from unisensory to multisensory areas. The modulation of the strength between synapses was performed given the co-occurrence of pre- and post-synaptic firing (i.e., Hebbian learning). The neural network was capable of accounting for the rapid recalibration of PPS over a wide range of time-constant values (from 2 to ∼7.5), while PPS was not plastic (on a rapid time-scale) in a smaller range (from 0 to 2), and vanished entirely for values beyond ∼7.5 (see Noel et al., 2018b, for a demonstration that the network’s capacity to engender a PPS effect is immune to changes in other parameter values). This pattern suggests that to the extent that rapid recalibration of PPS is scaffolded on a Hebbian mechanism, its rapid recalibration is not a peculiarity, but a piece carefully regulated to allow for plasticity and concurrent gradation between the near and far space. That is, a time constant of integration large enough that would not allow for rapid recalibration of PPS would also eliminate the representation of PPS altogether.

Importantly, we do not mean to suggest that Hebbian learning is the sole mechanism that could potentially explain rapid recalibration of PPS. For example, history-dependent modifications in the dynamical state of the network (i.e., a kind of working memory) could play an equally important role. For parsimony, however, we maintained the same mechanism used to model longer-term forms of PPS plasticity (i.e.: tool use, Serino 2015), and show that it can also account for rapid recalibration of PPS, when implemented in this more extreme, yet physiologically plausible form (synaptic plasticity has been observed to be effective even over timescales shorter than those modeled in our work; Arnsten et al. 2010). Finally, it is worth noticing that the time constant of the forgetting factor able to show recalibration was found to be the same order of magnitude of the time separating two consecutive multisensory stimulations. Future studies, both experimental and theoretical, could be performed to investigate the effect of increasing or decreasing the frequency of stimuli presentation. Moreover, additional models can be implemented that combine the short-term plasticity of the current model with the long-term synaptic maturation rules used in previous works (Magosso et al., 2010a; Serino et al., 2015a), to investigate the interplay of different timescale in PPS plasticity.

To conclude, here we show at the behavioral, electrophysiological, and computational level, a recalibration of the extent of PPS based on the type of multisensory interaction just experienced, on a trial-by-trial basis. This almost on-line regulation of the PPS boundary fits well with current accounts of PPS (Serino, 2019; Bufacchi & Iannetti, 2018; Clery et al., 2015; Brozzoli et al., 2014), which agree in describing it as multisensory-motor system mediating defensive and approaching interactions between the body and external stimuli. This space, therefore, needs to rapidly adapt to sudden changes in the environment to allow for adaptive responses.

## Contributions

J.P.N. designed research, performed research, analyzed data, and wrote the paper. T.B. performed research, analyzed data, and wrote the paper. E.T. performed research. E.P. performed research. B.H. contributed tools. C.C. contributed tools. E.M. performed research, contributed tools, and wrote the paper. O.B. contributed tools and wrote the paper. M.W. contributed tools and wrote the paper. A.S. designed research, contributed tools, and wrote the paper.

## Acknowledgments

The authors thank Jacob Feldman for help with figures. This study was supported by a grant from the Swiss National Foundation [PP00P3_163951/1], and by the Fondation Biaggi (Lausanne, Switzerland) to AS, by the Bertarelli Foundation and the Swiss National Science Foundation [Grant number 320030_166643] to O.B, and by NIH CA183492 and HD083211 to MTW. J.P.N. was additionally supported by NIH/NIMH [Grant number 1F31MH112336].

## Supplementary Figures

**Figure S1.**
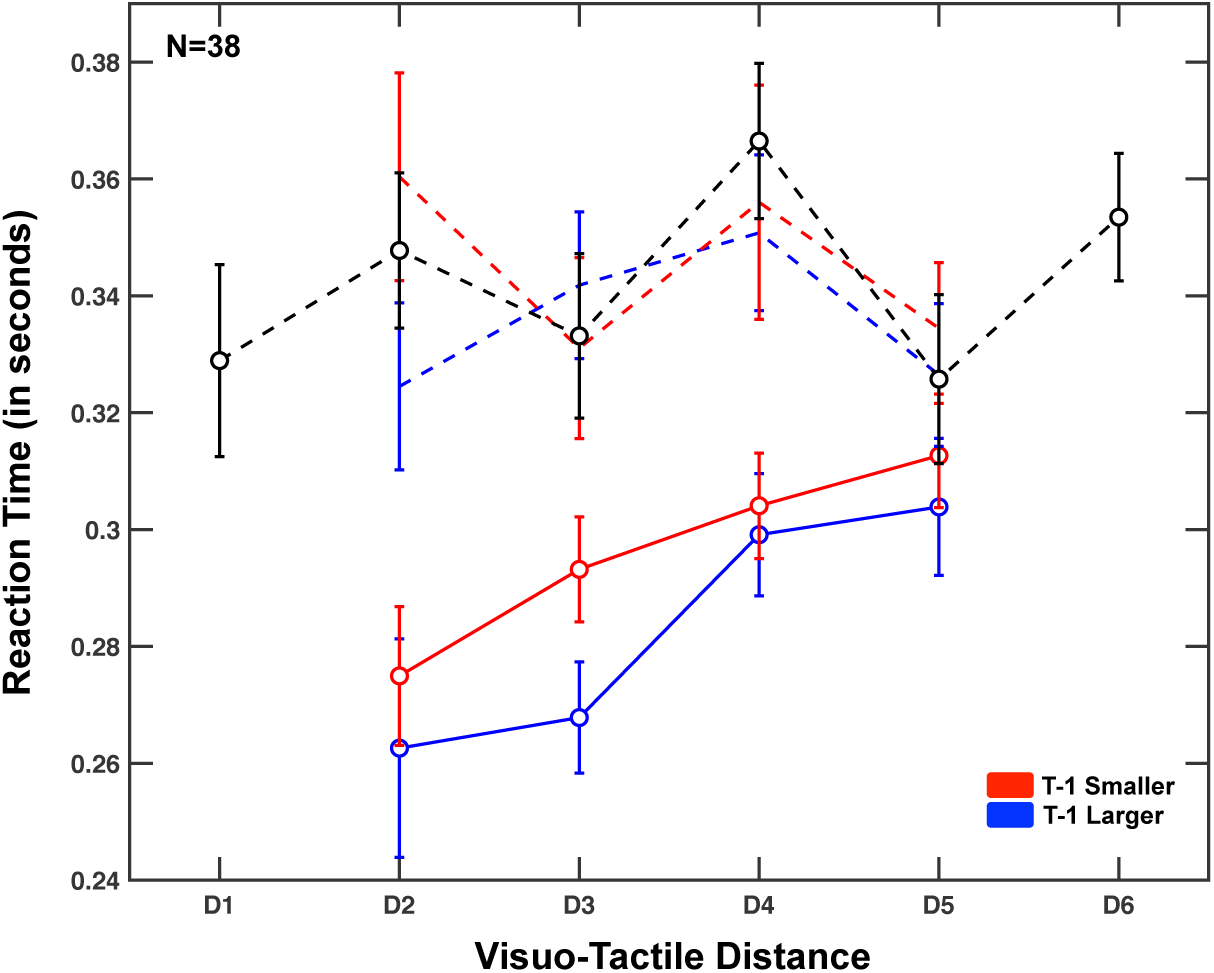
Rapid Recalibration of PPS, including all subjects. Given that a larger number of participants were removed from analyses in the main text given poor fits, we repeated the analyses while including all participants and performing simple analyses of variance, as opposed to data fitting. As in the main text, these analyses demonstrated a general visuo-tactile multisensory facilitation with respect to tactile reaction time (dashed vs. solid lines: t = 6.71, p = 7.03×10^11^). Further, a one-way ANOVA demonstrated that visuo-tactile proximity played a role in further enhancing multisensory facilitation (F = 5.04, p = 0.02). Most importantly, a 2 (t-1 smaller vs. t-1 larger) x 4 (distances: D2-D5), demonstrated a significant interaction (F = 3.37, p = 0.04), confirming that trial history impacts PPS encoding.

**Figure S2.**
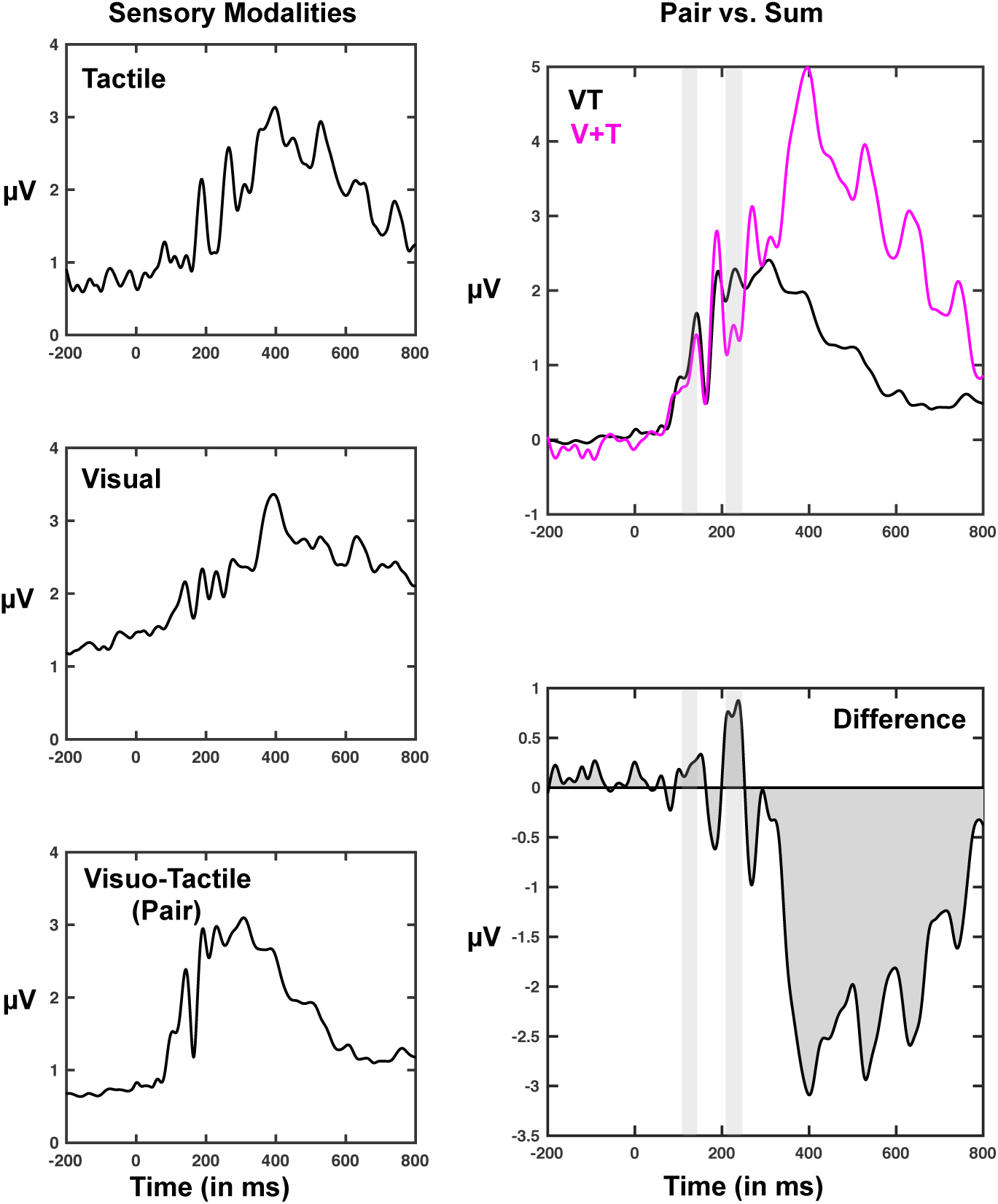
Global Field Power while subsampling trials to match across conditions. The results are virtually identical to those in the main text, strongly suggesting that given the number of repetitions represented we had reached an asymptote in signal-to-noise ratios. While matching the number of V, T, and VT repetitions at the single subject level, we observer supra-additivity (p < 0.01) between 123-160ms and 204-226ms post-stimuli onset.

**Figure S3.**
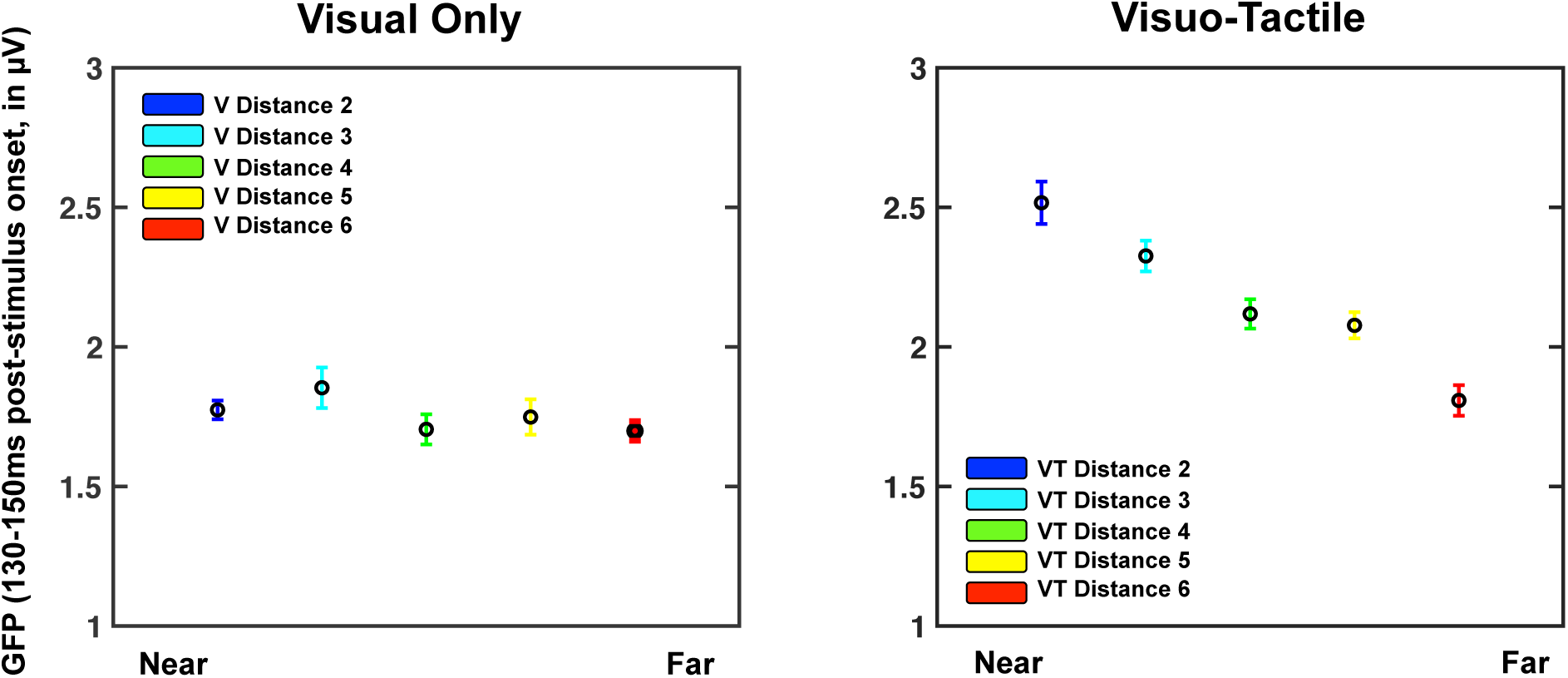
The modulation of GFP response as a function of distance is specific to VT stimuli. A 2 (V vs. VT) × 5 (distances) on the mean GFP between 130 and 150ms post-stimuli onset demonstrated a significant main effect of distance (F = 6.41, p = 6.15×10^−5^) and stimuli condition (F = 67.24, p = 1.11×10^−14^). Most importantly, there was a significant interaction between these variables (F = 28.79, p = 8.12×10^−20^) driven by the fact that while this peak was modulated by distance in VT presentations (main text), it was not in the case of visual-only stimulation (F = 1.17, p = 0.32).

